# Bacteroides thetaiotaomicron enhances H_2_S production in Bilophila wadsworthia

**DOI:** 10.1101/2024.10.14.618174

**Authors:** Jade Davies, Melinda J. Mayer, Nathalie Juge, Arjan Narbad, Lizbeth Sayavedra

## Abstract

Sulfate- and sulfite-reducing bacteria (SRB) are a group of strict anaerobes found within the human gut. *Bilophila wadsworthia*, a sulfite-reducing bacterium which produces hydrogen sulfide (H_2_S) from taurine and isethionate respiration is a common member of the healthy commensal human gut microbiota, but has been implicated in several disease states including inflammatory bowel disease and colorectal cancer. *Bacteroides thetaiotaomicron*, one of the most prominent gut bacteria, has sulfatases which release sulfate, serving as a potential substrate for sulfate-reducing bacteria. Here, we showed that when *B. thetaiotaomicron* and *B. wadsworthia* were in co-culture, there was a significant increase in *B. thetaiotaomicron*’s growth and in H_2_S production by *B. wadsworthia*. Differential gene expression analysis revealed increased expression of *B. wadsworthia*’s *dsrMKJOP* complex in co-culture, which delivers electrons for sulfite reduction to H_2_S. This was accompanied by a decreased expression of genes associated with taurine, sulfolactate and thiosulfate respiration, indicating that *B. thetaiotaomicron* may provide an alternative source of sulfite to *B. wadsworthia*. We hypothesised adenosine 5’-phosphosulfate (APS) to be this intermediate. Indeed, *B. wadsworthia* was able to grow using APS or sulfite as electron acceptors. Endometabolomic and transcriptomic analyses revealed decreased production of indole by *B. thetaiotaomicron* in co-culture with *B. wadsworthia* due to enhanced tryptophan utilisation by *B. wadsworthia*. The results of this microbe-microbe interaction could have significant pro-inflammatory effects in the human gut environment.

## Introduction

Sulfate- and sulfite-reducing bacteria (SRB) comprise a group of strict anaerobes and are found within the colonic mucosa as part of the commensal gut microbiota in at least 50% of humans ^1, 2^. SRB utilise inorganic sulfate (SO_4_^2–^) or sulfite (SO_3_^2–^) as a terminal electron acceptor during energy metabolism, a process which occurs concomitantly with oxidation of molecular hydrogen or organic compounds. Hydrogen sulfide (H_2_S) is the final product of dissimilatory sulfate reduction by SRB ^3^ and can freely diffuse across cell membranes ^4^. H_2_S is recognised as a hazardous product as it is both corrosive and toxic to many organs even at low concentrations ^5^. Classical sulfate-reducing bacteria reduce sulfate to sulfite via sulfate adenylyltransferase (Sat) and adenylylsulfate reductase (AprAB) ^6^ (Figure S1). Sulfite then enters the dissimilatory sulfite reduction pathway. *Bilophila wadsworthia* is a Gram-negative member of the SRB, first identified in 1989 from gangrenous and perforated appendicitis samples ^8^.Unlike other SRB, *B. wadsworthia* cannot utilise sulfate ^7–9^, and instead degrades organosulfate compounds such as taurine, isethionate, and sulfoquinovase, generating H_2_S as a by-product ^10, 11^. In contrast, the majority of living organisms have the assimilatory sulfate reduction pathway ^12^. This involves the reduction of sulfate to APS via sulfate adenylyltransferase (Sat), which is then converted to phosphoadenosine phosphosulfate (PAPS) via adenylylsulfate kinase / APS kinase. PAPS is then reduced to sulfite by phosphoadenosine phosphosulfate (PAPS) reductase, and the sulfite is then converted to H_2_S by assimilatory sulfite reductase ^12^ (Figure S1).

*B. wadsworthia* is present in the faeces of 50-60% of healthy individuals ^13^, and can also be isolated from buccal and vaginal samples ^14^. *B. wadsworthia* is considered to be virulent, as it is the third most common anaerobic isolate from appendicitis samples and appears to be clinically important in a variety of anaerobic infections ^15^. Furthermore, it exhibits endotoxic activity ^16^, and is adherent to human embryonic intestinal cells *in vitro* ^17^. Association studies in humans have linked *B. wadsworthia* enrichment in the gut with many diseases including colorectal cancer ^18^, multiple sclerosis ^19^, Parkinson’s disease ^20^, dementia ^21^, non-alcoholic steatohepatitis ^22^, intrahepatic cholestasis in pregnancy ^23^, diabetic kidney disease ^24^ and schizophrenia ^25^. Additionally, it was demonstrated that a human stool-derived *B. wadsworthia* strain was able to induce systemic inflammation in specific-pathogen-free mice ^26^. Given *B. wadsworthia*’s status as a potential pathobiont in the human gut, it is important to investigate the factors that influence its abundance and function.

*Bacteroides thetaiotaomicron* is a Gram-negative obligate anaerobe within the Bacteroidaceae family ^27^. Originally isolated from the faeces of a healthy adult ^27^, *B. thetaiotaomicron* is highly abundant constituting 1-6% of the total bacteria ^28^ with a 46% prevalence ^29^, and is a key commensal member of the human gut microbiota. *B. thetaiotaomicron* plays a major role in the human gut, including modulation of the host mucosal immune system ^30^, and utilisation of a wide range of polysaccharides ^31^. Some of these polysaccharides can be host-derived, and include glycosaminoglycans such as chondroitin sulfate, mucin, hyaluronate and heparan sulfate ^32^. The degradation of polysaccharides yields simple sugars for fermentation, resulting in the production of beneficial short-chain organic acids including acetate, lactate, succinate and propionate ^33^; *B. thetaiotaomicron* has also been shown to support the growth of butyrate-producing *Anaerostipes caccae in vitro* ^34^. Short-chain fatty acids (SCFAs) are a valuable source of energy for human colonocytes and aid in maintaining healthy barrier function ^35^. *B. thetaiotaomicron* encodes 28 sulfatases which can cleave sulfated residues of the host glycosaminoglycans to yield free sulfate in the mouse gut ^36^. By increasing the availability of free sulfate *in vivo*, *B. thetaiotaomicron* sulfatase activity could permit increased sulfate reduction and H_2_S production by SRB ^37^. Indeed, *B. thetaiotaomicron* and *Desulfovibrio piger,* a SRB, can co-colonise the gut, with *B. thetaiotaomicron* providing free sulfate to *D. piger* and promoting H_2_S production both *in vitro* and *in vivo* ^2^. Additionally, *B. thetaiotaomicron* generates hydrogen during polysaccharide fermentation and molecular hydrogen, which is used by hydrogenotrophs, including SRB, as an electron donor ^38, 39^. Given these established interactions with other SRBs, we aimed to further investigate the dynamics between *B. thetaiotaomicron* and *B. wadsworthia*. Here, we used transcriptomics and metabolomics to explore the mechanisms underpinning the interaction of *B. wadsworthia* and *B. thetaiotaomicron* during anaerobic co-culture.

## Results and Discussion

### Co-culture of *B. wadsworthia* and *B. thetaiotaomicron* boosts H_2_S production

*B. wadsworthia* (QI0013) and *B. thetaiotaomicron* (QI0072) were grown anaerobically in monocultures and co-cultures on synthetic (BPM, see methods) media supplemented with 10 mM taurine at a 1:1 ratio (10^6^ colony forming units (CFU)/mL inoculation density). At 8 hours post-inoculation, a significant increase in H_2_S concentration was observed in the co-abundance remained unaffected by the co-culture (Figure 1b). The elevated H_2_S levels were attributed to a significantly increased H_2_S production per *B. wadsworthia* cell when in co-culture with *B. thetaiotaomicron* (Figure 1d). Interestingly, the interaction also appeared to benefit *B. thetaiotaomicron*, which grew to significantly higher abundance in the co-culture (Figure 1c).

**Figure 1:**
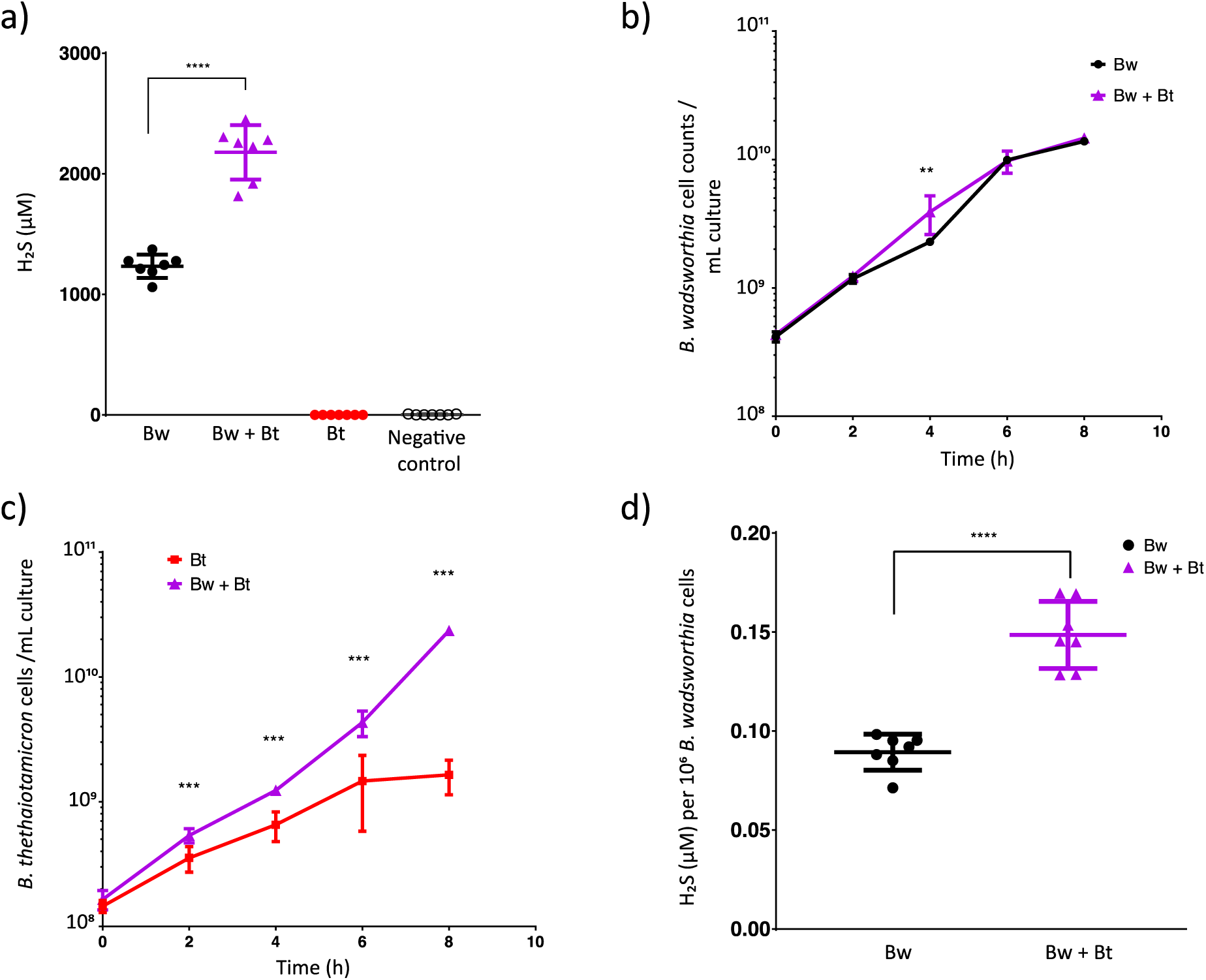
Co-culture of *B. wadsworthia* (Bw) QI0013 and *B. thetaiotaomicron* (Bt) QI0072. a) H_2_S concentrations (µM) at 8 h. b-c) Cell counts of *B. wadsworthia* or *B. thetaiotaomicron* during culture measured via qPCR. d) H_2_S concentration (µM) per 10^6^ *B. wadsworthia* cells at 8 h. Each point represents a culture replicate (n=7). In the negative control bacterial cells were not added. Horizontal lines represent average, and error bars represent SD. Results of unpaired t-tests are shown where ** = p ≤ 0.01, *** = p ≤ 0.001, **** = p ≤ 0.0001.

To gain further insights into the potential mechanism that increases H_2_S production in *B. wadsworthia*, we performed transcriptomic and metabolomic analysis of the mono and co-cultures at 8 hours. The most impacted pathways are discussed below.

### Respiration of sulfite via hydrogen or lactate by *B. wadsworthia* was increased in co-culture

In the co-culture, four genes associated with sulfur metabolism leading to H_2_S production in *B. wadsworthia* were overexpressed (Figure 2a, Table S1). These genes are part of the dissimilatory sulfite reductase protein complex (*dsrMKJOP*), including *dsrM* (WCP94_001671), *dsrK* (WCP94_001670), *dsrO* (WCP94_001668), and *dsrP* (WCP94_001667) (Table S1). When *B. wadsworthia* respires taurine, sulfite (SO_3_^2-^) generated enters the dissimilatory sulfite reduction pathway and is further reduced by DsrAB to produce H_2_S (Figure S1). Expression of *dsrMKJOP* can be increased under H_2_-rich growth conditions in *D. vulgaris* ^40^; the observed up-regulation in co-culture may suggest a higher bioavailability of H_2_ to *B. wadsworthia*. *B. thetaiotaomicron* can generate hydrogen (H_2_) during polysaccharide fermentation ^41^; therefore, *B. wadsworthia* could utilise this directly as an electron donor during dissimilatory sulfite reduction ^38, 42^. The increased expression of *dsrMKOP* in co-culture alludes to sulfite utilisation as an electron acceptor, potentially leading to increased H_2_S production.

**Figure 2:**
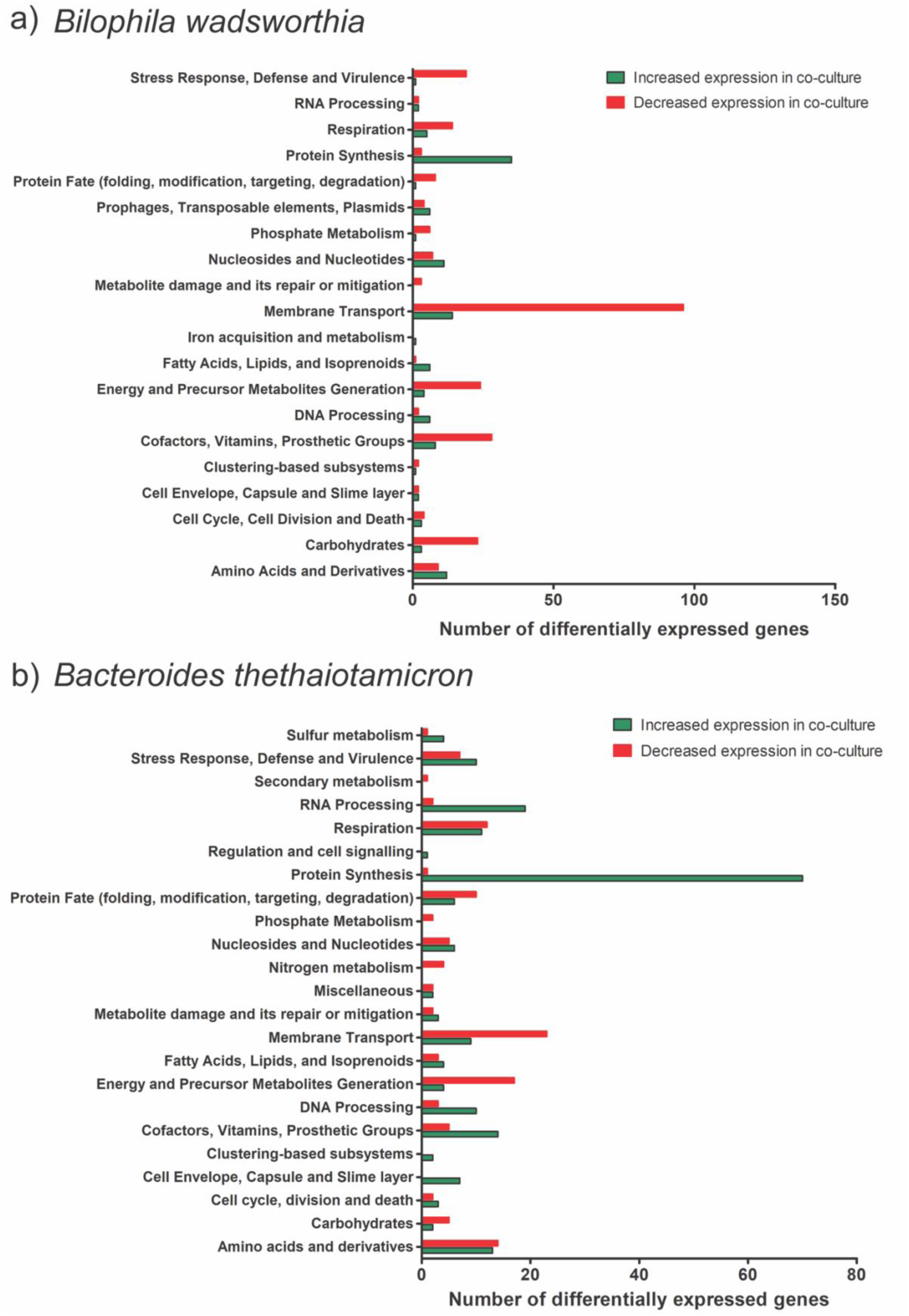
Differentially expressed genes (DEGs) of *B. wadsworthia* and *B. thetaiotaomicron* in co-culture versus monoculture. a) DEGs in *B. wadsworthia*. b) DEGs in *B. thetaiotaomicron*. Bar charts show numbers of DEGs increased (green) and decreased (red) in expression in co-culture relative to the respective monoculture in each functional gene class as annotated in BV-BRC.

In addition to hydrogen, lactate can also be utilised as an electron donor for the respiration of sulfite ^10, 43, 44^ (Figure 3). Interestingly, two genes encoding lactate permeases were overexpressed in co-culture (WCP94_002306, WCP94_000357) in *B. wadsworthia* (Table S1), suggesting enhanced utilisation of lactate. The presence of different electron donors might contribute to the increased expression of *dsrMKJOP* in *B. wadsworthia* under co-culture conditions, leading to a higher H_2_S production as a by-product. Taken together, the transcriptomic data revealed an increased capacity for uptake and utilisation of hydrogen and lactate in *B. wadsworthia* in co-culture with *B. thetaiotaomicron*; both compounds are key electron donors for dissimilatory sulfite reduction. This is in line with increased H_2_S concentration by *B. wadsworthia* in co-culture (Figure 1a).

**Figure 3:**
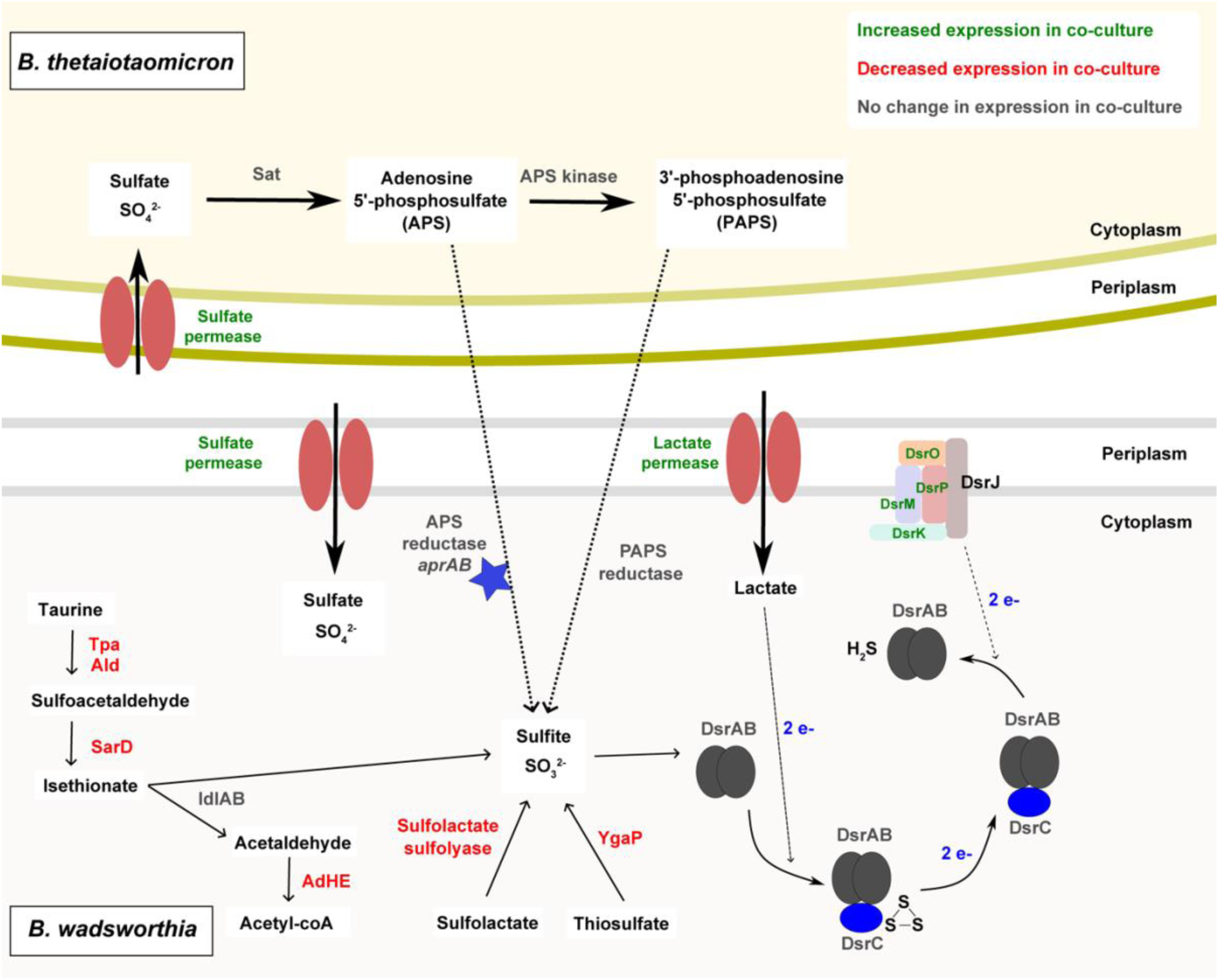
Model of cross-feeding interaction between *B. wadsworthia* and *B. thetaiotaomicron*. Enzymes are shown in colours corresponding to transcriptomic data: green showed increased gene expression in co-culture; red showed decreased expression in co-culture; grey showed no change in expression between co-culture and monoculture. The blue star represents a putative enzyme where functionality is unproven. The dotted line represents a putative cross-feeding mechanism. Dashed lines represent electron transfer.

Surprisingly, the gene cluster associated with taurine metabolism exhibited reduced expression in the co-culture, despite the observed increase in H_2_S concentration. This gene cluster includes taurine pyruvate dehydrogenase (*tpa*, WCP94_00949), alanine dehydrogenase (*ald*, WCP94_00950) and sulfoacetaldehyde reductase (*sarD*, WCP94_00947) ^10^ (Figure S1, Figure 3, Table S1). It has been shown that expression of these enzymes is elevated in *B. wadsworthia* cells grown in taurine, but not in isethionate-grown cells ^10^. Isethionate sulfite-lyase *islA* and isethionate sulfite-lyase activating protein *isiB* were not among the differentially expressed genes, but acetaldehyde dehydrogenase (*adhE*, WCP94_001941), which encodes an enzyme able to convert acetaldehyde (a product of isethionate degradation) to acetyl coA was decreased in expression in co-culture (Table S1). The downregulation of *tpa* and *sarD,* combined with the increased expression of *dsrMKJOP* suggests that *B. wadsworthia* can also utilise the respiration pathway from sulfite or a precursor molecule in the presence of *B. thetaiotaomicron* (Figure 3). Sulfite could be produced through different metabolic pathways in *B. wadsworthia*: i) from sulfolactate, via the activity of a sulfo-lyase (WCP94_000771-772) which converts sulfolactate to sulfite and pyruvate; this enzyme showed decreased expression in co-culture; or ii) through the inner membrane protein YgaP (WCP94_002882), a rhodanese-domain-containing protein that generates sulfite from thiosulfate, and which also showed decreased expression in co-culture (Table S1). Therefore, it is likely that in co-culture with *B. thetaiotaomicron*, *B. wadsworthia* utilises an alternative source of sulfite (Figure 3).

### Modulation of sulfur metabolism in *B. wadsworthia* and *B. thetaiotaomicron* co-culture

As taurine utilisation by *B. wadsworthia* as a sulfite source in co-culture appeared to be decreased, alongside an increased expression of genes associated with the dissimilatory sulfite reductase pathway and elevated H_2_S concentration, we investigated other possible sources of sulfite for *B. wadsworthia* in the co-culture. Increased expression of sulfate permease (WCP94_001830) and sulfatase (WCP94_001848) genes was observed in *B. wadsworthia* in co-culture (Table S1), suggesting an increased bioavailability of sulfate; however, whether these transporters are promiscuous for other compounds such as sulfite is unknown.

Given that *B. wadsworthia* cannot utilise sulfate, it was previously assumed that the genes for sulfate utilisation were absent from the genome ^7, 8, 45^. The *B. wadsworthia* QI0013 genome does not contain genes encoding the Sat enzyme necessary for sulfate reduction. However, our analysis revealed that *B. wadsworthia* QI0013 encodes two gene clusters of the alpha and beta subunits of the adenylylsulfate reductase (WCP94_000309-310 and WCP94_000741-742) (Table S3, Figure S1). Despite low amino acid sequence similarity to AprAB proteins from *Desulfovibrio* strains (Figure S2, Table S4), we identified conserved protein domains, including FAD-dependent oxidoreductase 2 and succinate dehydrogenase/fumarate reductase flavoprotein, supporting the prediction of AprAB activity (Table S5). Furthermore, we found that the putative AprAB genes were widely distributed in publicly available *B. wadsworthia* genomes (Table S6). This suggests a potential role for *B. wadsworthia* in APS reduction via adenylylsulfate reductase. The putative proteins encoded by these genes could facilitate the conversion of APS to sulfite, offering an alternative precursor molecule for the dissimilatory sulfite reduction pathway.

Interestingly, the *B. thetaiotaomicron* QI0072 genome encodes some enzymes involved in assimilatory sulfite reduction, including sulfate adenylyltransferase (Sat) (QI0072_1554, QI0072_1555) and adenylylsulfate kinase / APS kinase (QI0072_1556) (Table S3), which would catalyse the conversion of sulfate to both APS and PAPS. However, *B. thetaiotaomicron* QI0072 does not encode the enzymes for further utilisation of PAPS and APS. *B. wadsworthia* QI0013 expresses both phosphoadenosine phosphosulfate (PAPS) reductase (WCP94_00194), which reduces PAPS to sulfite, in addition to the putative AprAB which reduces APS to sulfite (Table S3).

We hypothesise that *B. thetaiotaomicron* QI0072 produces sulfate from sulfated saccharides via sulfatases ^46^, an activity which has been previously described to support the growth of *D. piger* via cross-feeding ^47^, and then reduces the sulfate to APS and/or PAPS. In *B. thetaiotaomicron*, genes encoding sulfate adenylyltransferase subunits 1 and 2 and adenylylsulfate kinase were consistently expressed at similar levels in both co-culture and monoculture (Table S3), suggesting the production of APS and PAPS. *B. thetaiotaomicron*-derived APS and PAPS could then be utilised directly by *B. wadsworthia*, where either AprAB or PAPS reductase yield sulfite, which then enters the dissimilatory sulfate reduction pathway (Figure 3). In this way, *B. thetaiotaomicron* may provide *B. wadsworthia* with an alternative source of sulfite, which may be energetically favourable over taurine degradation, or utilised when taurine or other organosulfur compounds are depleted. Indeed, we observed a decreased expression of genes associated with sulfite generation via taurine, thiosulfate and sulfolactate under co-culture conditions, with the phenotypic observation of increased H_2_S concentration (Figure 1). The adenylylsulfate reductase and PAPS reductase genes were expressed in *B. wadsworthia* in both co-culture and monoculture, without significant differences in expression levels (Table S3). Next, we investigated if *B. wadsworthia* would grow using sulfite or APS as electron acceptors instead of taurine. *B. wadsworthia* was able to show a significantly higher growth in media supplemented with 4 mM sulfite, 10 mM APS, or 10 mM taurine, compared to only anaerobe basal broth (ABB) (Figure 4).

**Figure 4:**
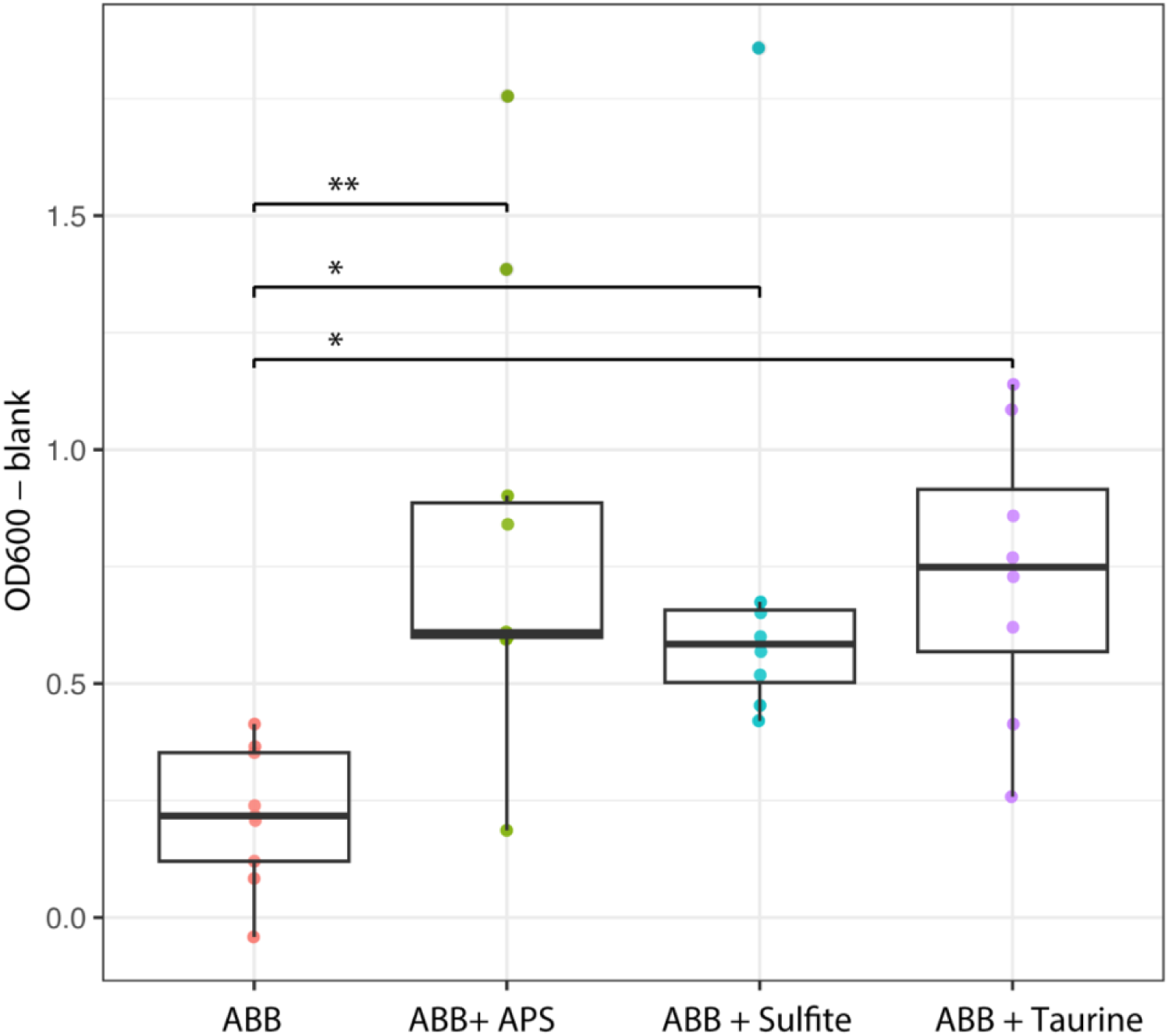
Growth (OD_600_) of *B. wadsworthia* in ABB media supplemented with 10 mM adenosine 5’-phosphosulfate (APS), 4 mM sulfite or 10 mM taurine. Growth was compared to *B. wadsworthia* grown in ABB media. Each box plot represents the median and interquartile range of the distribution of 7 culture replicate. Results of mixed linear model analyses are shown where * = p ≤ 0.01, ** = p ≤ 0.001.

Next, we tested the capacity of *B. thetaiotaomicron* to promote the sulfidogenic activity of other *B. wadsworthia* strains i.e *B. wadsworthia* QI0012, QI0014, and QI0015 strains. Co-culture with *B. thetaiotaomicron* resulted in significantly increased H_2_S concentration in all *B. wadsworthia* strains tested compared to their respective monocultures (Figure 5a), suggesting that the sulfidogenic potential of *B. thetaiotaomicron* is not strain specific. Indeed, QI0014 did not produce detectable H_2_S in monoculture at 8 h, whereas 195.3 ± 18.60 µM was produced in co-culture with *B. thetaiotaomicron* (Figure 5a). Interestingly, qPCR analysis of *B. wadsworthia* cell abundance revealed differences between strains with QI0012 and QI0013 showing slight increases in abundance in co-culture with *B. thetaiotaomicron* although this effect was not significant with QI0013 (Figure 5b), as shown above for this strain (Figure 1). A decreased abundance was observed in co-culture *B. wadsworthia* QI0014 and QI0015 strains with *B. thetaiotaomicron* (Figure 5b). Upon standardizing H₂S concentrations relative to *B. wadsworthia* cell numbers, it was observed that *Bacteroides thetaiotaomicron* consistently enhanced H₂S production across the various *B. wadsworthia* strains. Nonetheless, the specific mechanisms behind this increase may differ among strains (Figure 5).

**Figure 5:**
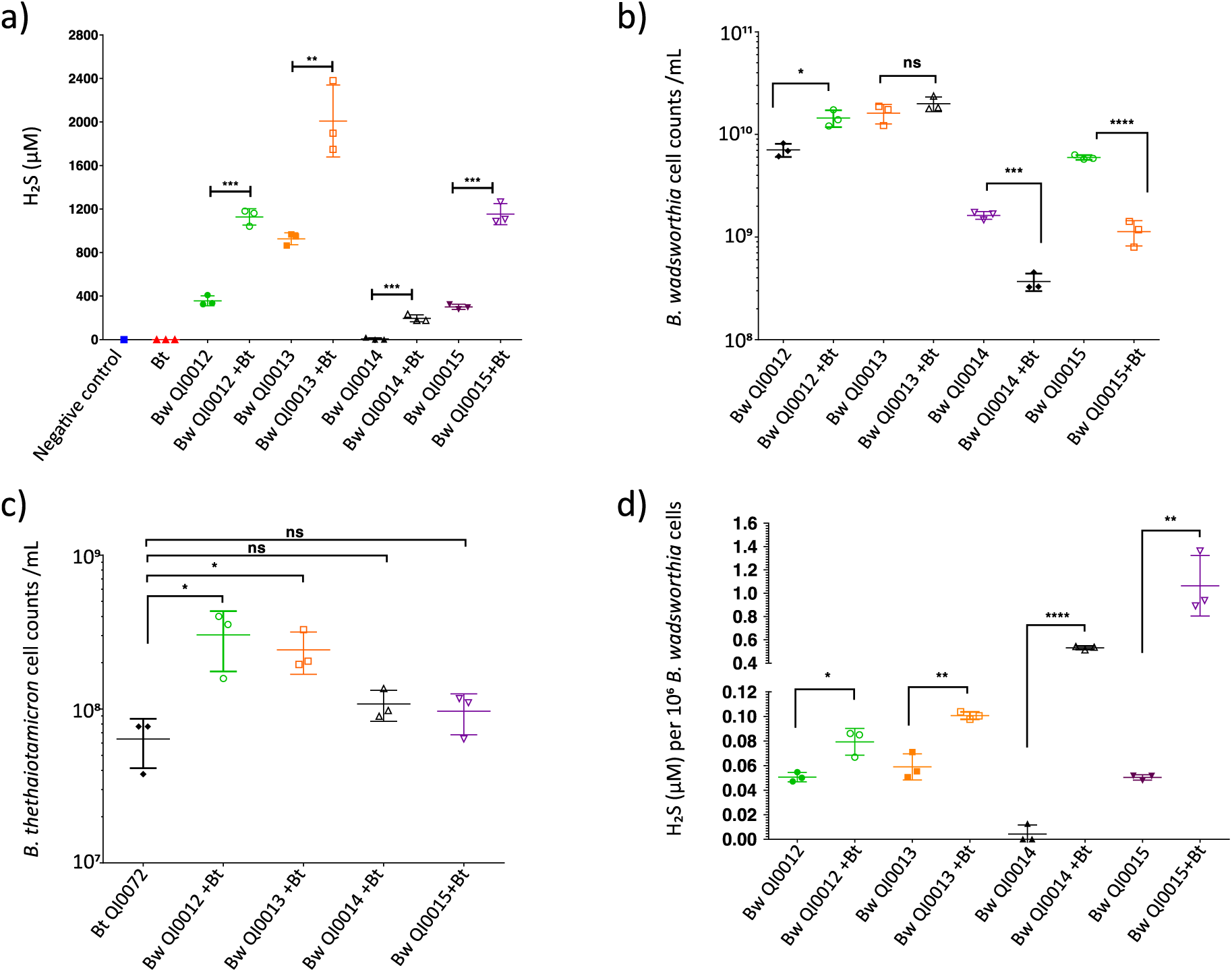
Pairwise co-culture of Bt strain 1 with four *B. wadsworthia* strains (QI0012, QI0013, QI0014, QI0015). a) H_2_S concentrations (µM) at 8 h. b) qPCR-determined *B. wadsworthia* cell counts at 8 h. c) qPCR-determined *B. thetaiotaomicron* cell counts at 8 h. d) H_2_S concentration (µM) per 10^6^ *B. wadsworthia* cells. Each point represents a culture replicate (n=3). Horizontal lines represent average, and error bars represent SD. Statistical significance between culture conditions was established using one-way analysis of variance (ANOVA) with Tukey’s multiple comparison tests with a significance level set at α = 0.05. Results show * = p ≤ 0.05, ** = p ≤ 0.01, *** = p ≤ 0.001, **** = p ≤ 0.0001, ns = not significant (p > 0.05).

The increased H₂S levels in QI0012 and QI0013 co-cultures are attributable to a combination of higher *B. wadsworthia* abundance and elevated H₂S production per cell. In contrast, the rise in H₂S observed with QI0014 and QI0015 is predominantly due to a substantial increase in H₂S production per *B. wadsworthia* cell (Figure 5d). Additionally, *B. thetaiotaomicron* exhibited a slight increase in abundance in all co-cultures compared to its monoculture (Figure 5c). Overall, *B. thetaiotaomicron* showed an ability to increase H_2_S production in all four *B. wadsworthia* strains that were tested.

### H_2_S-utilising amino acid biosynthetic pathways were reduced in expression in ***B. thetaiotaomicron***

Based on our genome analyses, *B. thetaiotaomicron* QI0072 does not encode known genes for H_2_S production, but it does have the capacity to utilise H_2_S during the biosynthesis of amino acids including cysteine and homocysteine. *B. thetaiotaomicron* QI0072 over-expressed twelve genes and under-expressed nine genes in co-culture associated with amino acid metabolism, compared to monoculture (Figure 2b). Interestingly, this strain under-expressed O-acetylserine sulfhydrolase (QI0072_3844) in co-culture (Table S2); this enzyme catalyses the conversion of O3-acetyl-L-serine and H_2_S to L-cysteine and acetate, utilising pyridoxal-5’-phosphate (vitamin B6) as a co-factor ^48^. This is relevant as this indicates that although *B. thetaiotaomicron* can utilise *B. wadsworthia*-derived H_2_S, the gene encoding this enzyme is heavily decreased in expression in co-culture with *B. wadsworthia*, which could indirectly contribute to the higher H_2_S concentration observed in the co-culture. Similarly, two genes involved in H_2_S-utilising homocysteine biosynthesis were also differentially expressed; one gene encoding O-acetylhomoserine sulfhydrylase (EC 2.5.1.49) was notably decreased in expression in co-culture (QI0072_3143, -5.12 logFC), whereas another gene encoding this enzyme was increased in expression in co-culture (QI0072_2658, 1.24 logFC) (Table S2). This enzyme catalyses the conversion of O-acylhomoserine and H_2_S to homocysteine ^49, 50^. Overall, decreased homocysteine and cysteine biosynthesis by *B. thetaiotaomicron* in co-culture with *B. wadsworthia* results in lower H_2_S utilisation by *B. thetaiotaomicron*, which indirectly contributes towards the overall phenotype of high H_2_S concentration in the co-culture.

### Alterations to the endometabolome in *B. wadsworthia* and *B. thetaiotaomicron* co-culture

The endometabolome of the *B. wadsworthia* and *B. thetaiotaomicron* co-culture was analysed at 8 hours post-inoculation, alongside monocultures. The *B. thetaiotaomicron* monoculture metabolite abundance clustered distinctly from the *B. wadsworthia* monoculture and co-culture conditions based on partial least squares discriminant analysis (PLS-DA) (Figure S3a). The heatmap displaying the top 50 differentially abundant metabolites showed clear differences between the conditions, most notably when the *B. thetaiotaomicron* monoculture is compared to the co-culture conditions (Figure S3b). The variable importance in projection (VIP) scores showed that the top 15 compounds contributing to differences between conditions were relatively high in abundance in the *B. thetaiotaomicron* monoculture and in relatively low abundance in the co-culture (Figure not produced by *B. thetaiotaomicron* in co-culture with *B. wadsworthia* or consumed by *B. wadsworthia*. The highest-scoring compound was Ala-Val, followed by the ester phenethyl acetate, sphinganine and nicotinic compound 1-Methylnicotinamide (Figure 6a). Kynurenic acid also had a significantly lower abundance in co-culture compared to *B. thetaiotaomicron* monoculture (Figure 6a,b), suggesting a reduction in tryptophan metabolism via this pathway. Kynurenic acid is a product of tryptophan degradation in mammals ^51^, however, *Escherichia coli* has been demonstrated to produce kynurenic acid from L-kynurenine in the rat small intestine ^52^ and the human gut microbiota may also produce this compound, since kynurenic acid concentration is high in the distal colon relative to other body sites ^53^. Sphinganine can be produced by *Bacteroides* strains in the gut and has the potential to control the levels of bioactive lipids in the liver ^54^, underscoring its potential role in host-microbial interactions.

**Figure 6:**
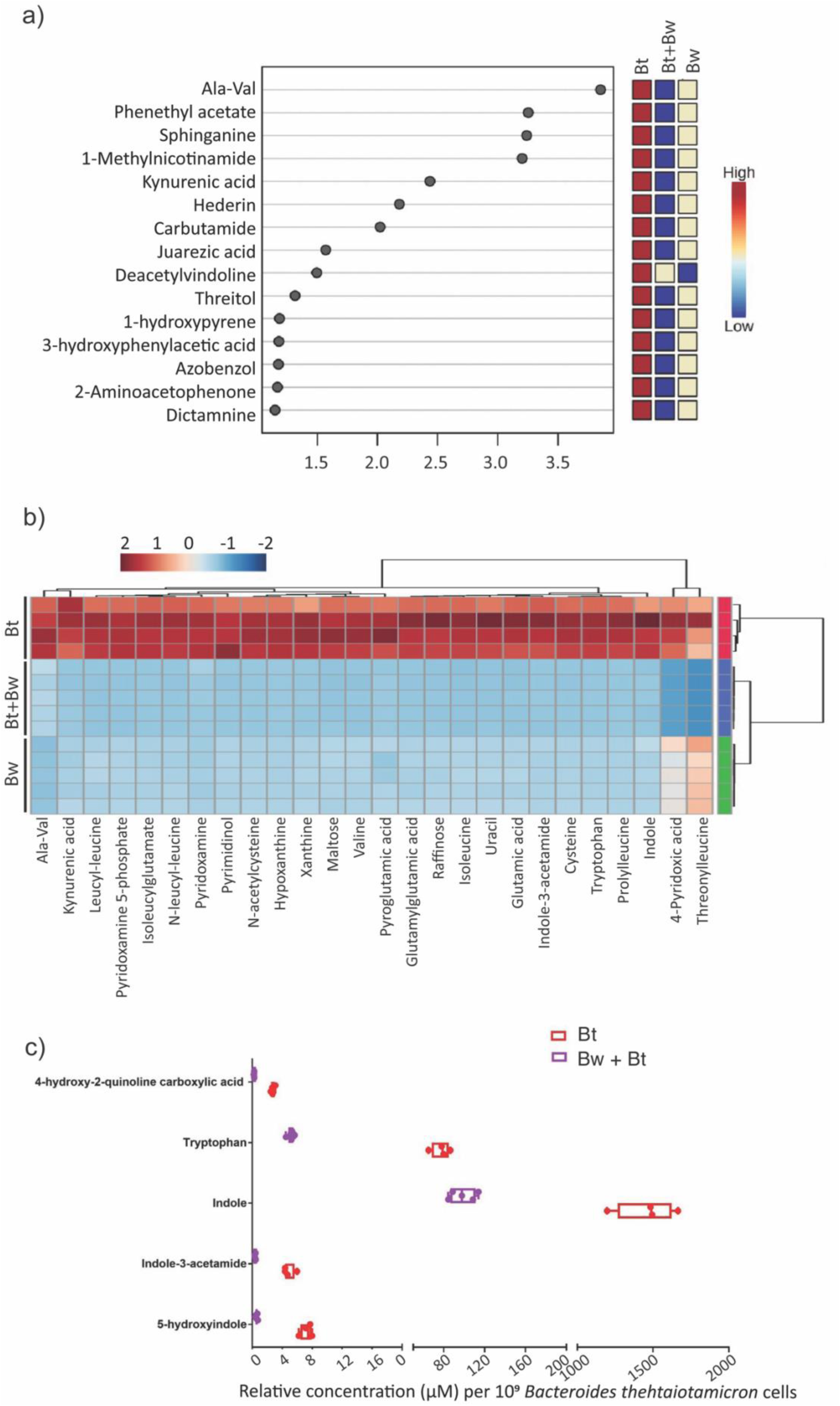
Comparisons of the endometabolome of *B. wadsworthia* and *B. thetaiotaomicron* in co-culture (Bw + Bt) with monocultures (Bw_mono, Bt_mono). a) The top compounds ranked based on the Variable Importance in Projection (VIP) scores. The coloured boxes on the right indicate the relative concentrations of the corresponding metabolite in each group. b) Specific metabolites of interest manually curated from the compound list detected via untargeted LC-MS. c) Relative intracellular concentrations of compounds related to tryptophan metabolism in *B. thetaiotaomicron* (Bt) in monoculture and co-culture with *B. wadsworthia* (Bw + Bt). Concentration is standardised to *B. thetaiotaomicron* cell counts. Each point represents a culture replicate. Box and whisker plots show line at mean, box at 25^th^ – 75^th^ percentile, whiskers to minimum and maximum.

The transcriptomic data identified branched-chain amino acids (BCAAs) as significant metabolites of interest, showcasing differential expression in Liv system genes responsible for ABC transporters specific to BCAAs in *B. wadsworthia* (Figure 2a, supplementary Table S1). This may signify a modified bioavailability of leucine, isoleucine, and valine for *B. wadsworthia* in the co-culture environment. Indeed, valine and isoleucine were detected in the endometabolome in all culture conditions, in addition to dipeptides of leucine including threonylleucine, prolylleucine, N-leucyl-leucine and leucyl-leucine (Figure 6b). In all cases, relative abundance was high in *B. thetaiotaomicron* monoculture, lower in *B. wadsworthia* monoculture and the lowest abundance was in the co-culture condition (Figure 6b). Taken together, this suggests lower bioavailability of BCAAs in the endometabolome of *B. wadsworthia* and *B. thetaiotaomicron* in co-culture at 8 hours compared to monocultures. This interplay could have implications for human health, since increased concentrations of aromatic and BCAAs by the gut microbiome have been linked to insulin resistance and type 2 diabetes mellitus ^55^.

### Impact on tryptophan metabolism between *B. thetaiotaomicron* and *B. wadsworthia*: insights from combined metabolomic and transcriptomic analyses

The endometabolome revealed differences in the relative concentrations of tryptophan, indole, and indole-3-acetamide between *B. thetaiotaomicron* and *B. wadsworthia* monocultures and their co-culture; the abundance of these metabolites was higher in the *B. thetaiotaomicron* monoculture compared to the other culture conditions (Figure 6b and 6c). Bacterial produced indole has been shown to modulate intestinal inflammation in animals, and inhibit quorum sensing and virulence factor production ^56^. Indole-producing members of the human gut microbiota include *E. coli*, *Proteus vulgaris*, *Paracolobactrum coliforme*, *Achromobacter liquefaciens* and *Bacteroides spp*. ^57^. These strains can degrade tryptophan to indole, pyruvate and ammonia via the enzyme tryptophanase (TnaA), the activity of which is induced by tryptophan ^58^ and inhibited by glucose ^57^. Interestingly, indole production from tryptophan is commonly observed in *B. thetaiotaomicron* ^59^, and *B. thetaiotaomicron*-derived indole has been shown to inhibit virulence in enteropathogenic *E. coli* (EPEC) and *Vibrio cholerae* by inhibiting T3SS expression ^59^. Given that *B. wadsworthia* cannot produce indole ^60^, it is likely that indole and its by-products are derived from *B. thetaiotaomicron*. In co-culture, the relative abundance of these metabolites decreased compared to the *B. thetaiotaomicron* monoculture (Figure 6b). To confirm that this effect was not due to the concentrations being standardised to the total cell count, the indole and tryptophan relative abundances were determined in the *B. thetaiotaomicron* monoculture and co-culture, normalized to 10^9^ *B. thetaiotaomicron* cells only as determined by qPCR. This showed a decreased relative abundance of indole and tryptophan in the endometabolome of the co-culture compared to *B. thetaiotaomicron* monoculture (Figure 6c). This suggests that *B. wadsworthia* may either utilise indole derived from *B. thetaiotaomicron* or directly uses tryptophan, thereby limiting the availability of tryptophan for *B. thetaiotaomicron*. Transcriptomic data supported this observation, showing increased expression of a tryptophan-specific transport protein (WCP94_002980) and tryptophanyl-tRNA synthetase gene (WCP94_002967) in *B. wadsworthia* during co-culture, indicating enhanced tryptophan uptake and incorporation into new proteins. These combined metabolomic and transcriptomic findings suggest that, under co-culture conditions, *B. wadsworthia* intensifies its tryptophan uptake and protein incorporation, potentially diminishing tryptophan availability for *B. thetaiotaomicron*’s indole and kynurenic acid production. These observations underscore a dynamic interplay between the two bacteria in the utilisation of tryptophan and its downstream metabolites in the co-culture environment.

## Conclusions

The increased H_2_S production by *B. wadsworthia* when co-cultured with *B. thetaiotaomicron* could have implications for human gut health. Both H_2_S and *B. wadsworthia* have been implicated in gut inflammation and pathogenesis, making it crucial to understand the change in H₂S production. Our co-culture experiments demonstrated relationship. Transcriptomic analysis of *B. wadsworthia* revealed reduced expression of genes involved in sulfite production from taurine, thiosulfate, and sulfolactate, suggesting an alternative sulfite source in the co-culture. Our integrated transcriptomic and genomic data indicate a putative cross-feeding mechanism, with *B. thetaiotaomicron* potentially providing APS to *B. wadsworthia* for sulfite production; we next demonstrated that *B. wadsworthia* can grow to high densities using APS as an electron acceptor, substantiating this proposed cross-feeding interaction. Gene expression data suggested an increased uptake of lactate in *B. wadsworthia*, which could be used as an electron donor in dissimilatory sulfite reduction. Beyond sulfur metabolism, we observed alterations in tryptophan metabolism in co-culture. *B. wadsworthia*’s increased expression of genes associated with tryptophan uptake suggests a competition for tryptophan, potentially impacting *B. thetaiotaomicron*’s indole production. Reduced indole production by *B. thetaiotaomicron* in co-culture aligns with increased tryptophan utilisation by *B. wadsworthia*, hinting at potential implications for gut health and inflammation ^61, 62^. In conclusion, the increased H_2_S production by *B. wadsworthia* in co-culture, coupled with the reduced H_2_S utilisation by *B. thetaiotaomicron*, underscores a complex microbial interaction. This proposed APS-mediated cross-feeding mechanism opens avenues for further exploration into sulfate metabolism in the human gut. The overall shift towards decreased indole and increased H_2_S production in co-culture suggests potential pro-inflammatory effects in the human gut environment, warranting further investigation.

## Materials and Methods

### B. thetaiotaomicron and B. wadsworthia isolation and growth

*B. thetaiotaomicron* QI0072 was isolated from a faecal sample donated by a healthy adult between 50-80 years old recruited via the COMBAT study (ClinicalTrials.gov Identifier: NCT03679533). An initial enrichment was obtained in anaerobic anaerobe basal broth (ABB) liquid broth media (Oxoid, Thermo Fisher Scientific) supplemented with 10 mM taurine, and the isolate was purified on Brain Heart Infusion agar (BHI, Oxoid) prepared according to the manufacturer’s instructions then supplemented with 0.5% v/v vitamin K solution in ethanol (10 µL/L), resazurin (1 mg/L), hemin (5 mg/L) and L-cysteine hydrochloride (0.5 g/L) (further referred as BHI+C) with 1.5% (w/v) agar. Additional chemicals were from Merck. *B. wadsworthia* strains QI0012, QI0013, QI0014 and QI0015 were isolated from stool samples from healthy human donors recruited via the QIB Colon Model study (ClinicalTrials.gov Identifier: NCT02653001) using ABB supplemented with 10 mM taurine as described by Sayavedra *et al*. ^63^. Cell pellets were cryopreserved at -80°C using Protect Select Anaerobe Cryopreservation tubes (Technical Service Consultants, UK).

### Bacterial co-culture experiments

BHI, de Man, Rogosa and Sharpe (MRS, Oxoid) and ABB liquid broth media were prepared according to the manufacturer’s instructions. BHI + supplements (BHI+S) was prepared by adding hemin (10 mg/L) and yeast extract (5 g/L) to BHI broth before autoclaving. Where required, solid media was made by the addition of 1.5% (w/v) agar to the media before autoclaving. All media and culture vessels were maintained under anaerobic conditions using an anaerobic cabinet (Don Whitley, UK) with materials pre-reduced before use for at least 18 hours in an atmosphere of 5% CO_2_, 10% H_2_ in N_2_ at 37°C. For co-culture experiments, *B. thetaiotaomicron* was grown on BHI+C, and *B. wadsworthia* was grown on ABB supplemented with 10 mM taurine. After overnight incubation, the second passage cultures were diluted to ∼10^8^ CFU/mL using estimated cell count per optical density at 600 nm (OD_600_) factor. OD_600_ was quantified using a SPECTROstar Nano instrument (BMG Labtech). Co-culture experiments used anaerobic synthetic media containing a mix of 1:1:1 (v/v) of BHI+C, MRS and Postgate C, supplemented with 10 mM taurine (further referred to as BPM media). Postgate C contained per litre of distilled water: sodium lactate (6 g), sodium sulphate (4.5 g), ammonium chloride (1 g), yeast extract (1 g) potassium dihydrogen phosphate (0.5 g), sodium citrate tri basic (0.3 g), magnesium sulphate 7-hydrate (0.06 g), iron sulphate 7-hydrate (4 mg), calcium chloride (0.04 g), L-cysteine hydrochloride (0.5 g), and resazurin (0.8 mg) ^64^.

Co-cultures were prepared by inoculating ∼10^6^ CFU/mL anaerobic BPM. The CFU used for inoculum were confirmed by plating on ABB supplemented with 10 mM taurine for *B. wadsworthia*, and BHI+C media for *B. thetaiotaomicron*. Experimental conditions included negative control (no inoculum), monocultures and co-culture with final culture volumes of 10 mL. All cultures were performed in at least triplicate. Sub-samples were taken at 0, 2, 4, 6 and 8 h post-inoculation.

### Growth of *B. wadsworthia* with different electron acceptors

The growth of *B. wadsworthia* under various conditions was assessed by monitoring its OD_600_ after 48 hours of incubation on ABB media enriched with different electron phosphosulfate (APS) sodium salt (Merck, UK), and 10 mM taurine, alongside a control containing ABB only. We used at least seven technical replicates. Differences in growth were tested using a linear model with the glmmTMB package.

### Colourimetric determination of H_2_S concentration

Determination of H_2_S concentration in samples was performed using the methylene blue assay modified from Cline ^65^. Briefly, 500 µL of bacterial cultures were taken and immediately fixed 1:1 with 5% zinc acetate and stored at -20°C. For calibration, zinc sulfide solutions were prepared in media diluted 1:200 within concentration range 0 – 40 µM. For analysis, fixed samples were diluted 1:100 in water to a final volume of 1 mL, producing a final sample dilution of 1:200. 80 µL of diamine reagent (250 mL 6M HCl, 1g N,N-dimethyl-1,4-phenylendiamine sulfate, 1.5 g FeCl_3_.6H_2_0) was added to all samples and standards which were then stored in the dark to allow methylene blue colour development. After 30 min, the samples were centrifuged at 13,000 x g for 5 min to pellet biomass. 300 µL of supernatant was taken for spectrophotometric absorbance measurement at 670 nm. H_2_S concentration in µM was determined using the calibration curve as a standard. Statistical significance between culture conditions was established using unpaired t-test pair-wise comparisons and displayed graphically using GraphPad Prism 7 (GraphPad Software, Boston, USA). Results of unpaired t tests are shown, where * = p ≤ 0.05, ** = p ≤ 0.01, *** = p ≤ 0.001, **** = p ≤ 0.0001, ns = not significant (p > 0.05).

### Absolute quantification of bacterial cells via qPCR

For quantification of bacterial cells in experimental cultures, DNA extraction was performed using the Maxwell® RSC Blood DNA kit (Promega) according to the manufacturer’s protocol. Prior to extraction, 200 µL samples were boiled at 90°C for 10 min and 150 µL was transferred to a sterile microcentrifuge tube containing 30 µL of proteinase K solution and 300 µL lysis buffer. Each sample was vortexed for 10 s and incubated at 56°C for 20 min. Quantification of *B. wadsworthia* and *B. thetaiotaomicron* was performed via qPCR using KiCqStart® SYBR® Green qPCR ReadyMix™ with ROX (Sigma-Aldrich) on a StepOnePlus™ Real-Time PCR System (Thermo Fisher Scientific). Absolute quantification of gene copy numbers was performed by comparing samples to calibration standards prepared with known gene copy numbers in the range of 10 – 1 x 10^9^ copies. The number of gene copies per cell was used to calculate the absolute cell counts per mL of culture. All samples and standards were assayed in triplicate. The reaction conditions and primers used are shown in Table 1. Statistical significance between culture conditions was established using unpaired t-test pair-wise comparisons and displayed graphically using GraphPad Prism 7.

**Table 1:**
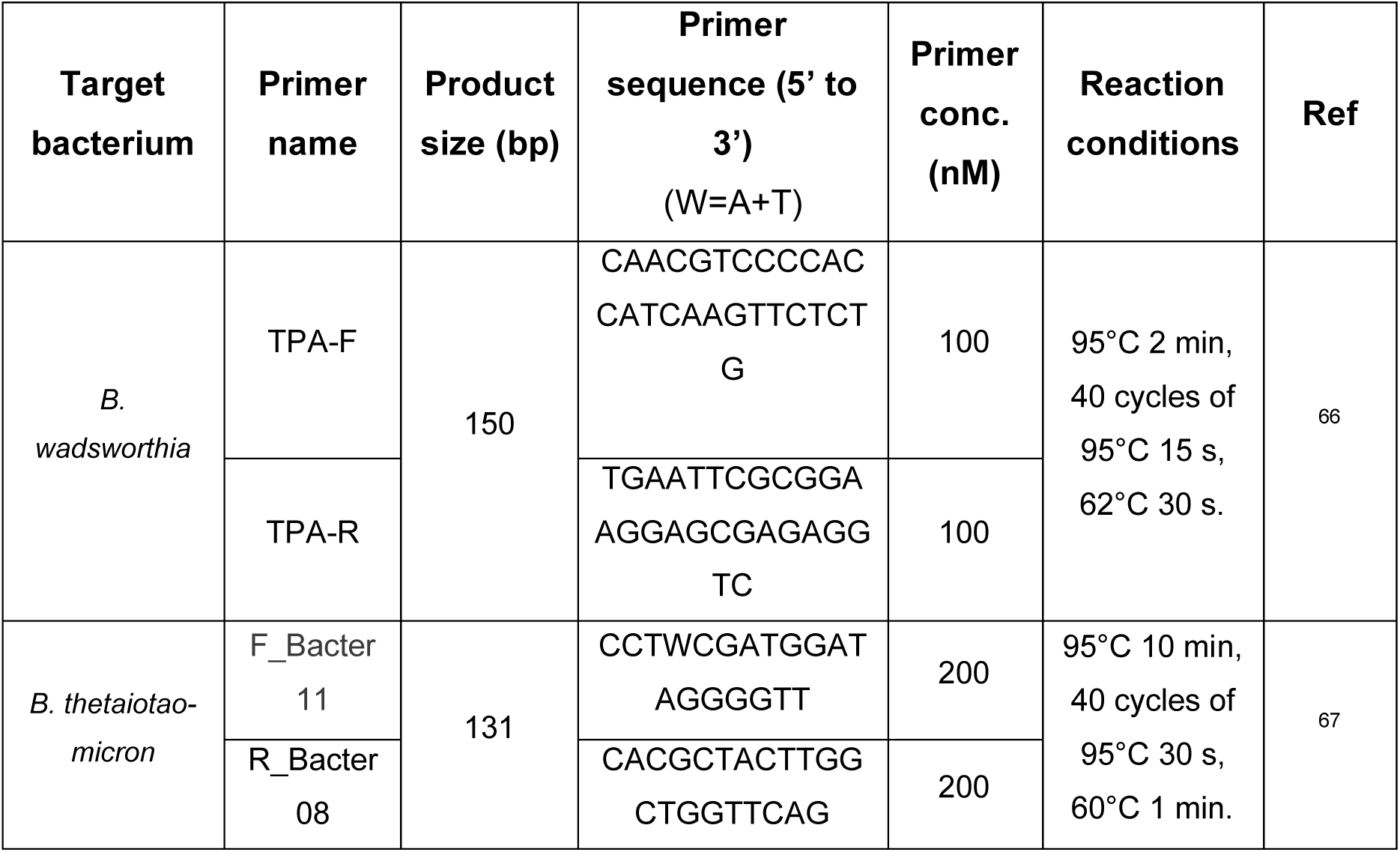
Primers and reaction conditions used for absolute quantification of *B. wadsworthia* and *B. thetaiotaomicron* via qPCR.

### Transcriptomic sequencing and analysis

5 mL of cultures were taken at 8 h post-inoculation and cells were immediately pelleted at 15,000 x g for 2 min. The supernatant was removed, cell pellets were snap frozen on dry ice, and stored at -80°C. RNA was extracted using the RNeasy® Mini Kit (Qiagen, Germany) according to manufacturer’s protocol with on-column DNase digestion. RNA concentration was determined via Qubit RNA High Sensitivity kit (Thermo Fisher Scientific, Massachusetts, USA) and Qubit 3.0 fluorometer (Life Technologies, Massachusetts, USA). Total RNA was sequenced at Azenta Genewiz. rRNA was depleted using NEBNext rRNA depletion kits for human/mouse/rat and bacteria species (E6310 and E7850). Samples were sequenced using an Illumina NovaSeq 6000 instrument with 2×150 bp configuration using a sequencing depth of 20 M paired-end reads per sample.

For differential expression analysis, raw data was analysed as previously described ^68^. Briefly, reads were cleaned to remove sequencing adapters and residual ribosomal RNA sequences using BBDuk (v 38.06, Bushnell B. - sourceforge.net/projects/bbmap/). Cleaned reads were mapped to the QI0013 and QI0072 reference genomes using BBSplit (v 38.06). The number of transcripts per gene was estimated using featureCounts (v.2.0)^69^. Reference genomes were annotated with the BV-BRC comprehensive genome analysis tool ^70^. Differentially expressed genes were identified using edgeR with TMM normalisation ^71^ using the Trinity RNASeq package ^72^, with a P value cut-off of 0.01 and a log2FC of 1 equating to a 2-fold change in gene expression. In the case of specific genes of interest where there was no functional annotation, InterProScan ^73^ was used to assign predicted protein functions based on domains where possible using the predicted amino acid sequence. Additionally, Pathway Tools v23.0 ^74^ was used for preliminary pathway enrichment analysis and pathway predictions using the reference genomes.

### Endometabolome analysis via LC-MS

5 mL of cultures were taken at 8 h post-inoculation and cells were immediately pelleted at 4,000 x g for 10 min. All supernatant was removed, and cell pellets were snap-frozen on dry ice. All LC-MS analysis was performed by Creative Proteomics, New York, USA. Prior to analysis, bacterial pellets were thawed and 240 µL methanol added for metabolite extraction. Samples were vortexed for 60 s, sonicated for 30 min at 4°C and stored at - 20°C for 1 h. Samples were pelleted at 12,000 x g for 15 min at 4°C. Finally, 200 μL of supernatant and 5 μL of DL-o-Chlorophenylalanine (0.2 mg/mL) was transferred to a vial for LC-MS analysis. QC samples were prepared by pooling all the samples in triplicate. All samples were injected in triplicate. Separation was performed by ACQUITY UPLC (Waters) combined with Q Exactive MS (Thermo) and screened with ESI-MS. The LC system was comprised of ACQUITY UPLC HSS T3 (100×2.1 mm×1.8 μm) with ACQUITY UPLC (Waters). The mobile phase was composed of solvent A (0.05% formic acid water) and solvent B (acetonitrile) with a gradient elution (0-1 min, 5% B; 1-12.5 min, 5%-95% B; 12.5-13.5 min, 95% B; 13.5-13.6 min, 95%-5% B; 13.6-16 min, 5% B). The flow rate of the mobile phase was 0.3 mL/min. The column temperature was maintained at 40°C, and the sample manager temperature set at 4°C. Mass spectrometry parameters in ESI+ and ESI-mode were as follows: ESI+: Heater Temp 300°C; Sheath Gas Flow rate, 45 arb; Aux Gas Flow Rate, 15 arb; Sweep Gas Flow Rate, 1 arb; spray voltage, 3.0 KV; Capillary Temp, 350°C; S-Lens RF Level, 30%. ESI-: Heater Temp 300°C, Sheath Gas Flow rate, 45 arb; Aux Gas Flow Rate, 15 arb; Sweep Gas Flow Rate, 1 arb; spray voltage, 3.2 KV; Capillary Temp, 350°C; S-Lens RF Level, 60%.

Metabolites were identified using Compound Discoverer 3.0 (Thermo Fisher Scientific). Progenesis QI v 2.1 (Waters) was used for manual screening of the identified compounds, in order to minimise false positive identification results. Data was normalised using the Total Ion Count (TIC) method where the peak area of each metabolite was divided by the SUM of all metabolites area and then multiplied by one million.

To remove compounds with high analytical variability, compounds with RSD_QC_ >20% were discarded ^75^. Endometabolomic data was standardised by calculating the concentration (µM) per 10^9^ bacterial cells present as measured via qPCR, in order to account for cell density differences in monocultures and co-cultures ^76^. Compounds at low concentrations across all samples (<2 µM per 10^9^ cells) were removed. In positive ion mode, 65% of the identified metabolites remained after quality control (320 left from 490), and in negative mode 70% remained (267 left from 377). Given the higher peak intensity and number of reported compounds, positive mode was used for further analysis. The metabolomic data was auto-scaled and analysed using Metaboanalyst 5.0 ^77^, to obtain the PLS-DA for the global profile changes and Variable Importance in Projection (VIP) compounds that contribute highly to inter-condition differences, and the heatmaps showing feature clustering and inter-condition differences in relative abundance of compounds.

### DNA extraction from isolates for whole genome sequencing

For whole genome sequencing, 1 mL of overnight culture was taken and centrifuged at 13,000 g for 2 min. Pellets were stored at -20°C until extraction. For extraction of high molecular weight DNA, Fire Monkey High Molecular Weight DNA (HMW-DNA) extraction kit was used according to manufacturer’s protocol (Revolugen, UK). The additional overnight elution step recommended by the manufacturer was used to maximise DNA yield. DNA was quantified using Qubit High Sensitivity DNA Quantification Kit (Thermo Scientific, UK) and Qubit 3.0 fluorometer.

### Whole genome sequencing

For whole genome sequencing of *B. thetaiotaomicron*, a hybrid approach using both short- and long-read sequencing was performed in-house at Quadram Institute Bioscience. For short-read sequencing, genomic DNA was normalised to 5 ng/µl with EB buffer (10 mM Tris-HCl) and sequencing was performed using an Illumina NextSeq 500 system with 2×150 bp paired-end reads. Libraries were prepared using Bead Linked Transposomes were received as fastq files. For long-read sequencing, a minION was used (Oxford Nanopore Technologies, Oxford, UK).

For genome assembly, short reads were cleaned using BBDuk (v 38.79) to trim the reads and remove sequencing adaptors. Long reads were trimmed using Porechop (v 0.2.3) ^78^ and a hybrid assembly was reconstructed using Unicycler (v 0.4.9) ^79^. The completeness and contamination of the assembly was checked using CheckM (v 1.0.18) ^80^. Genomes were annotated using Prokka (v 1.14.6) and BV-BRC ^81, 82^.

### Sequence comparison of sulfur metabolism genes

For genes of interest, amino acid sequences were obtained from *B. wadsworthia* (QI0013) and *B. thetaiotaomicron* strain 1 (QI0072) genomes in addition to those from reference genomes *D. desulfuricans* subsp. *desulfuricans* DSM 642, *D. gigas* DSM 1382 and *D. alaskensis* G20 via BV-BRC ^70^. InterProScan ^73^ was used to assign predicted protein function based on domains. Protein-protein comparison was performed using blastp on NCBI-BLAST using default settings ^83^. Amino acid alignments were performed using Clustal omega v1.2.2 in Geneious prime v 2022.1.1 (Biomatters Ltd) and visualised with GeneDoc v 2.7.000 (National Resource for Biomedical Supercomputing).

### Ethics statement

The colon model study was approved by the Quadram Institute Bioscience Human Research Governance Committee (IFR01/2015), and London - Westminster Research Ethics Committee (15/LO/2169). The COMBAT study was approved by the University of East Anglia’s Faculty of Medicine and Health Sciences Ethical Review Committee (Reference: 201819–039) and the Health Research Authority (IRAS number: 237251). The colon model study was registered under the ClinicalTrials.gov Identifier NCT02653001 and the COMBAT was registered under the NCT03679533.

## Acknowledgements

The authors thank David Baker (Quadram Institute Bioscience) for library preparation and whole genome sequencing.

## Data availability

Transcriptomic sequencing data was submitted to NCBI under PRJNA1115121. The genome of *Bilophila wadsworthia* QI0013 is available under project PRJNA1085689. The genome of *Bacteroides thetaiotaomicron* was submitted under project PRJNA1151643.

## Authors’ contributions

JD, LS and AN conceived and designed the project. JD and LS designed the experiments. JD performed experiments and analysed the data. LS supervised transcriptomic analyses. JD and LS wrote the manuscript with contributions from all co-authors.

## Supplementary data

**Figure S1:**
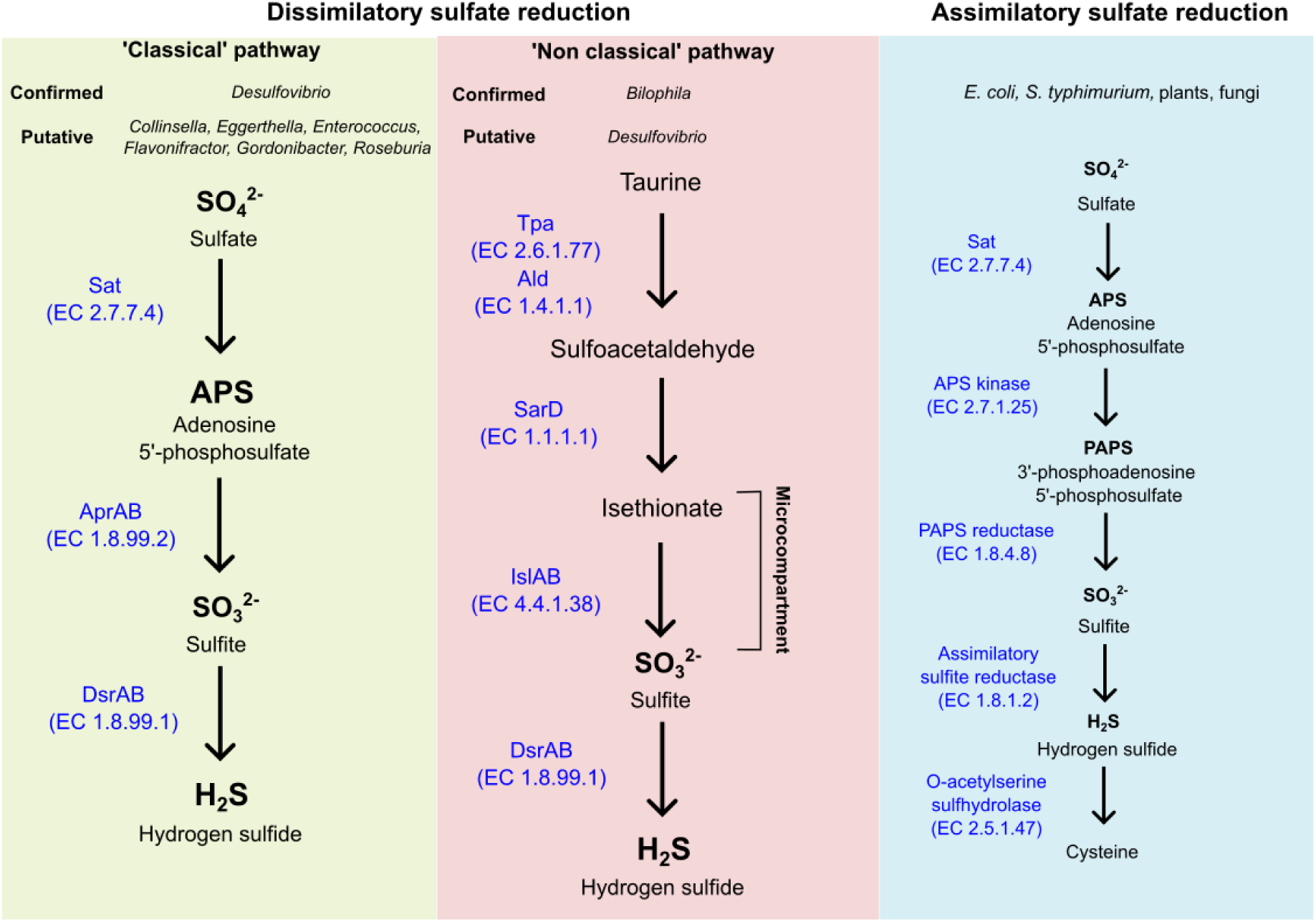
Metabolic pathways for sulfate reduction. Dissimilatory sulfate reduction is a strictly anaerobic process performed by sulfate-reducing bacteria (SRB). Toxic isethionate intermediates are encapsulated within microcompartments in *B. wadsworthia* ^1^.

**Figure S2:**
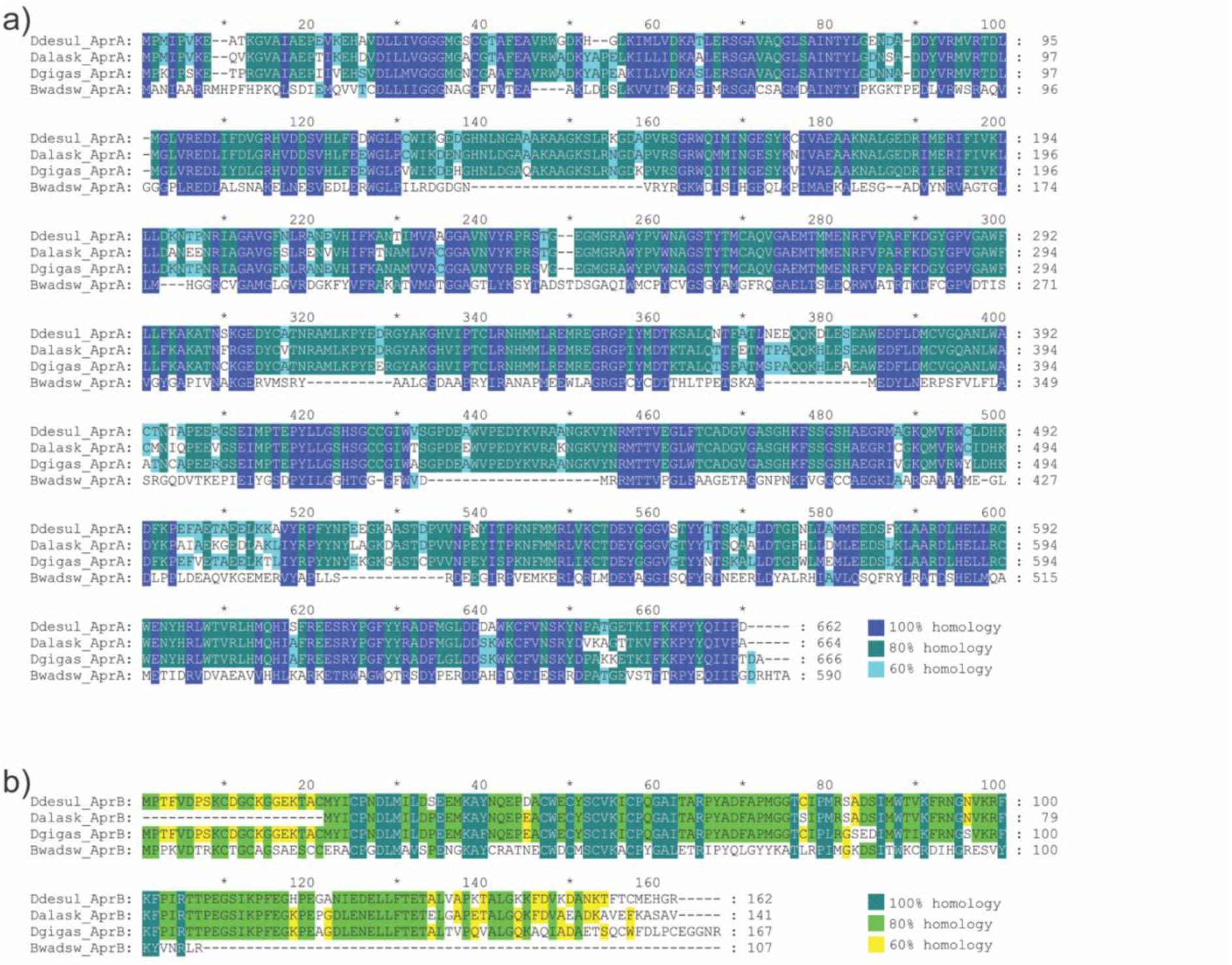
Alignment of amino acid sequences of A) AprA genes and B) AprB genes from *B. wadsworthia* QI0013 (Bwadsw), *Desulfovibrio gigas* DSM 1382 (Dgigas), *Desulfovibrio alaskensis* G20 (Dalask) and *Desulfovibrio desulfuricans* subsp. desulfuricans DSM 642 (Ddesul).

**Figure S3:**
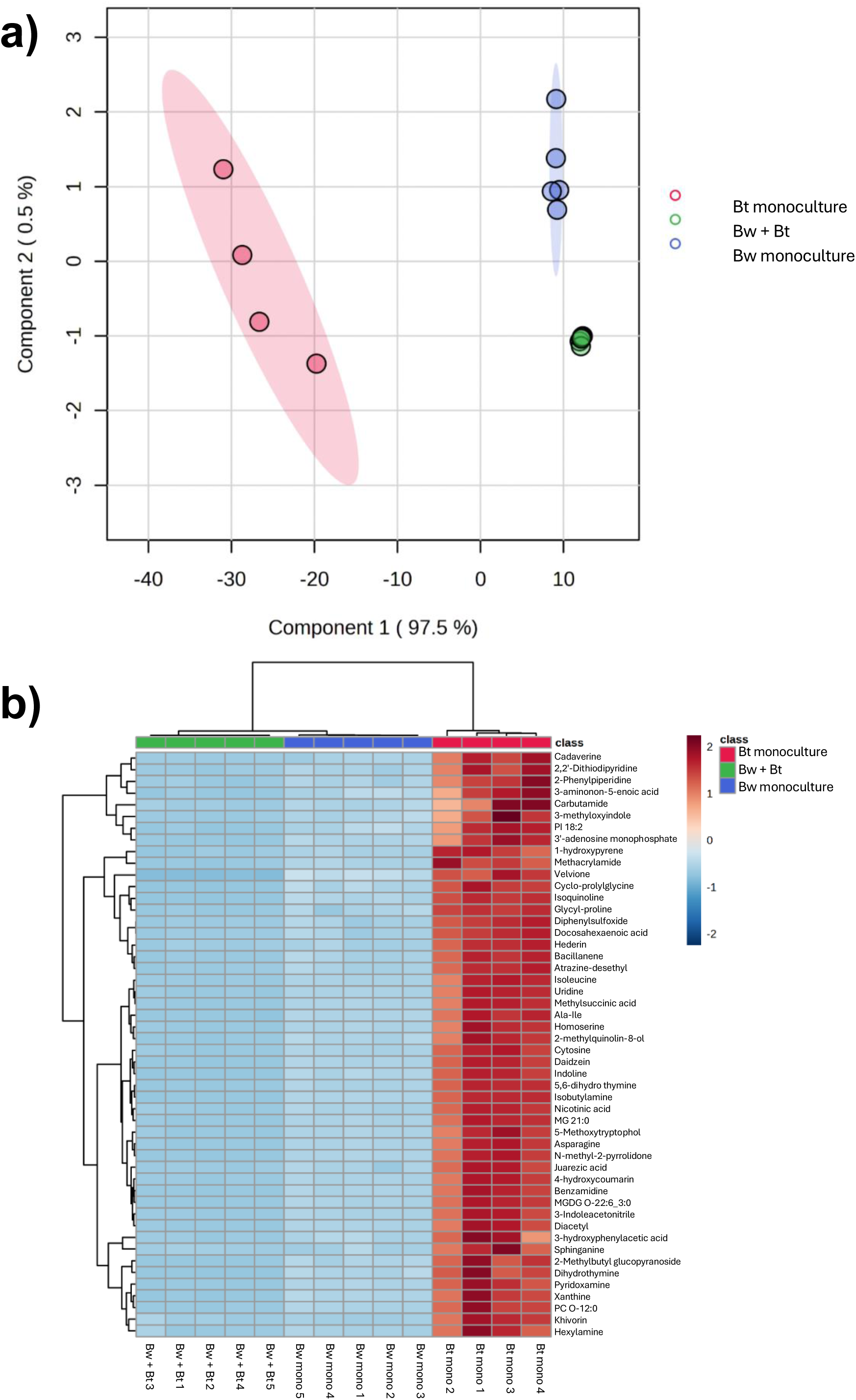
Comparisons of the endometabolome of *B. wadsworthia* and *B. thetaiotaomicron* in co-culture (Bw + Bt) with monocultures (Bw_mono, Bt_mono). a) PLS-DA plot of metabolites acquired from samples via untargeted LC-MS in positive ion mode. b) Heatmap displaying relative abundance of top 50 differentially abundant metabolites in the culture conditions.

